# Linking endoplasmic reticulum stress to polyploidy in ovarian cancer cells

**DOI:** 10.1101/2020.01.24.918029

**Authors:** Lucile Yart, Daniel Bastida-Ruiz, Christine Wuillemin, Pascale Ribaux, Mathilde Allard, Pierre-Yves Dietrich, Patrick Petignat, Marie Cohen

## Abstract

Polyploid giant cancer cells (PGCCs) have been observed in epithelial ovarian tumors and have the ability to survive to antimitotic drugs. Their appearance can result from paclitaxel treatment or hypoxia, two conditions known to induce unfolded protein response (UPR) activation. PGCCs produced under hypoxia may be formed by cell fusion and can contribute by bursting and budding to the generation of cancer stem-like cells which have a more aggressive phenotype than the parental cells. Despite the fact that PGCCs may contribute to tumor maintenance and recurrence, they were poorly studied. Here, we confirmed that PGCCs could derive, at least in part, from cell fusion. We also observed that PGCCs nuclei were able to fuse. The resulting cells were able to proliferate by mitosis and were more invasive than the parental cancer cells. Using two different ovarian cancer cell lines (COV318 and SKOV3), we showed that UPR activation with chemical inducers increased cell fusion and PGCCs appearance. Down-regulation of the UPR-associated protein PERK expression partially reversed the UPR-induced PGCCs formation, suggesting that the PERK arm of the UPR is involved in ovarian PGCCs onset.

## 1. Introduction

Epithelial ovarian cancer (EOC) is the most common type of ovarian cancer, representing 9 out of 10 tumors [1]. It is the seventh most common cancer in women and the leading cause of gynecologic cancer-related deaths in Western countries [2]. EOC is managed by cytoreductive surgery followed by platinum-based chemotherapy with an expected response rate to primary therapy of more than 70% [3]. Nevertheless, 2 years after a complete remission, most of the patients had a cancer recurrence [4] with often platinum-based chemotherapy treatment resistance [3].

EOC relapses origin is not completely understood, however, several hypotheses have been proposed over time. One of them, the cancer stem cell (CSC) theory, is based on the capacity of these cells to generate a proliferative progeny, initiating the growth of a heterogeneous tumor [5]. CSCs present chemotherapy resistance, allowing their survival to the treatment with subsequent proliferation and cancer relapse [6]. The role of CSCs in the ovarian cancer was reviewed by Giornelli et al., highlighting the importance of these cells in cancer recurrence and progression [7]. Despite the origin of CSCs remains elusive, some theories move towards a possible genesis in cancer cell fusion events [reviewed at [8]]. The fusion of cancer cells could give rise to polyploid giant cancer cells (PGCCs) [9], large cells containing several copies of DNA and positive for cancer stem cell markers [10]. Interestingly, the presence of PGCCs in ovarian cancer has been reported preferentially at late disease stages and in high pathological grades [10]. We previously showed that primary EOC cells isolated from malignant ascites were able to spontaneously form PGCCs in culture [11]. However, the origin of PGCCs is controversial since both fusion or endoreplication processes could give rise to them [8,9,12]. Zhang et al., demonstrated that in SKOV3, an EOC cell line established from ascites, the formation of PGCCs was at least partially due to cell fusion events [9]. In the study conducted by Zhang et al., the PGCCs derived from EOC cells had striking properties such as different cell cycle regulation and division profile, changing their daughter generation mechanism from mitosis to budding, which confer these cells the resistance to cytotoxic chemotherapy treatments [9]. Additionally, it was found that hypoxia increased PGCCs formation [9,13–15], but other conditions such as chemotherapy could also favor their production [13].

The tumor environment is known to be hypoxic and hypoglycemic due to the low irrigation caused by the fast and erroneous angiogenesis that tumor mass undergo [reviewed at [16]]. These conditions are capable of triggering the endoplasmic reticulum stress (ERS) response or unfolded protein response (UPR) which is essential for cellular adaptation to stressful situations [17]. In homeostatic conditions, the three UPR-associated proteins, protein kinase RNA-like endoplasmic reticulum kinase (PERK), inositol-requiring enzyme 1*α* (IRE1*α*) and activating transcription factor 6 (ATF6), remain inactive because of their binding with Glucose-regulated protein 78 (GRP78) [review at [18]]. Under ERS conditions, unfolded proteins accumulate in the ER lumen, promoting the recruitment of chaperones to enable their correct folding. Among these chaperones, GRP78 leaves the complexes formed with the UPR-associated proteins and induces the UPR activation [19]. This gives rise to a general decrease of protein, by contrast with increased expression of chaperones and ER-associated protein degradation-related proteins (ERAD) and autophagy activation [review at [18]]. The overexpression of GRP78 serves as UPR activation marker together with C/EBP homologous protein (CHOP) [20]. In ovarian cancer cells, GRP78 was shown to be overexpressed [21], suggesting UPR activation in these cells. Interestingly, GRP78 plays also an important role in trophoblastic cell fusion [22,23]. We thus hypothesized that UPR activation could be involved as well in EOC cell fusion and hence the occurrence of PGCCs.

In this article, we confirmed Zhang et al. results stating that PGCCs derive at least partially from cell fusion in SKOV3 cells [9]. Moreover, we observed that PGCCs nuclei are able to fuse under antibiotic selection pressure and we unraveled the advantages that nuclei fusion confers to these cells. Finally, we demonstrated that ERS induces cell fusion and PGCCs formation in EOC cells.

## 2. Materials and Methods

### 1.- Reagents

#### Chemical hypoxia

Cobalt Chloride (CoCl_2_, Merck, Darmstadt, Germany).

#### ER stress inducers

HA15 (Selleckchem, Zurich, Switzerland), Thapsigargin (THA, Enzo LifeSciences, Lausen, Switzerland) and Tunicamycin (TUN, Enzo LifeSciences, Lausen, Switzerland).

#### ER stress inhibitors

4-(2-aminoethyl) benzenesulfonyl fluoride hydrochloride (AEBSF; Sigma, Darmstadt, Germany), Melatonin (Mel; Sigma, Darmstadt, Germany), STF-083010 (STF; Selleckchem, Zurich, Switzerland), 4μ8C (Selleckchem, Zurich, Switzerland), GSK2656157 (GSK; Selleckchem, Zurich, Switzerland) and Salubrinal (SAL; Tocris Biosciences, Bristol, UK).

#### PERK silencing

PERK siRNA (h) (siPERK; SantaCruz Biotechnology,) and control siRNA-A (siControl; SantaCruz Biotechnology).

#### Chemotherapeutic drug

Paclitaxel (Ptx; Sigma, Darmstadt, Germany).

### 2.- Cancer cell line and culture

The human ovarian cancer cell lines COV318 (purchased from ECACC, via Sigma-Aldrich) and SKOV3 (courtesy of Dr Florence Delie, University of Geneva, Switzerland) were cultured at 37°C and 5% CO2 in DMEM (Gibco, Invitrogen, Basel, Switzerland) or RPMI (Gibco, Invitrogen, Basel, Switzerland) respectively, supplemented with 10% fetal bovine serum (FBS, Biochrom AG, Oxoid AG, Basel, Switzerland) and 0.05 mg/ml gentamycin (Invitrogen, Basel, Switzerland).

### 3.- Cell treatments

#### ERS induction

ERS was induced in ovarian cancer cells by adding either 100 nM THA, 1 μg/mL TUN or 1 μM HA15 in culture medium. Cells were treated for 48h before processing for analysis.

#### UPR pathways inhibition

UPR pathways were inhibited in ovarian cancer cells by adding either 17.5 μM SAL, 0.3 μM GSK, 8 μM 4μ8C, 2 μM STF, 200 μM AEBSF or 1 mM Mel, in combination with 100 nM THA or 1 μg/mL TUN. Cells were treated for 48h before processing for analysis.

#### PERK silencing

SKOV3 and COV318 cells were transfected either with 10 nM PERK siRNA or 10nM control siRNA using INTERFERin transfection reagent (Polyplus transfection, Illkirch, France). Twenty-four hours after transfection, ERS inducers were added in culture medium as described above.

### 4.- Generation of polyploid ovarian cancer cells by cell-cell fusion

Cytoplasmic staining and FACS sorting: SKOV3 or COV318 cells were stained with either green or far red cytoplasmic dyes, CellTrace CFSE (green; Life Technologies, Carlsbad, USA) and CellTrace Far Red (far red, Life Technologies, Carlsbad, USA), according to the manufacturer’s instructions. For each cell line, 3.5 × 10^5^ green cells and 3.5 × 10^5^ far red cells were co-seeded immediately after staining in a T175 flask. Twenty for hours after seeding, cells were treated with 300 μM CoCl_2_ for 48h. Cells were then analyzed by flow cytometry with FACSAria II (Becton Dickinson, Franklin Lakes, USA) for -GFP and far red signals. Double-positive cells were sorted and seeded in regular culture medium. Characteristic pictures of the sorted cells were done 16h after seeding, using fluorescence microscope (EVOS FL, Life Technologies, Carlsbad, USA).

Transfection and selection: SKOV3 cells were transfected with pH2B-mCherry-IRES-puro2 plasmid (Addgene, LGC Standards, Teddington, UK) or with pEGFP-CenpA-IRES-neo (Addgene, LGC Standards, Teddington, UK) using JetPEI as transfection reagent (Polyplus transfection, Illkirch, France) according to the manufacturer’s instructions. Two days after transfection, cells were selected using selection antibiotics (1 μg/ml puromycin (InvivoGen, San Diego, USA) for cells transfected with pH2B-mCherry-IRES-puro2 plasmid = SKOV3-Red, and 500 μg/ml G418 (Roche Life Science, Penzberg, Germany) for cells transfected with pEGFP-CenpA-IRES-neo = SKOV-3Green) for two weeks. Cells were then cultured in RPMI medium supplemented with 0.1 μg/ml puromycin for SKOV3-Red cells or with 50 μg/ml G418 for SKOV3-Green cells).

SKOV-M cell line establishment: SKOV3-Red and SKOV3-Green were mixed (1:1) and cultured for 24h before adding 0.2 μg/ml puromycin and 100 μg/ml G418 in RPMI. Culture medium was renewed every two days. Cell growth stabilized within three weeks.

### 5.- Cell cycle analysis

SKOV3-Green, SKOV3-Red and SKOV3-M (1 × 10^6^ of each) were resuspended in PBS with 5 μg/mL Hoechst 33342 (Life Technologies, Carlsbad, USA), and incubated for 20 min at 37°C in water-bath. Amounts of cells in G0-G1 and G2 phases were then assessed by flow cytometry analysis (LSR Fortessa, Becton Dickinson, Franklin Lakes, USA). Data were analyzed with FlowJo software (FlowJo LLC, Ashland, USA). This experiment was repeated 2 times.

### 6.- Cell proliferation

On day 0, SKOV3-Green, SKOV3-Red and SKOV3-M were resuspended in Hanks’ Balanced Salt Solution (Life Technologies, Carlsbad, USA) and incubated with 1 μM CellTrace Far Red (Life Technologies, Carlsbad, USA) for 7 min at 37°C. Staining was stopped by adding RPMI-10% FBS into cell suspension (1:1) for 5 min on ice. Cells were then washed twice with RPMI-10% FBS before seeding into 10 cm culture dishes (5 × 10^5^ cells/dish, 4 dishes/cell type) with regular culture medium. For each cell type, 5 × 10^5^ additional cells were immediately processed for far red signal analysis by flow cytometry (Gallios, Beckman Coulter, Brea, USA) (D0). Far red signal dilution was then daily measured for 4 days (D1, D2, D3 and D4). Data were analyzed with FlowJo software (FlowJo LLC, Ashland, USA). This experiment was repeated 3 times.

### 7.- Cell viability (3-(4,5-dimethylthiazol-2-yl)-2,5-diphenyltetrazolium bromide (MTT) assay)

Normalization of Invasion assay: SKOV3-Red, SKOV3-Green and SKOV3-M cells were seeded in 96-well plates at a density of 15’000 cells/well and incubated for 48h. Culture medium was then replaced by 100μl of MTT reagent. Plates were analyzed after 2 hours using a microplate reader at 540 and 690 nm (as blank). These experiments were carried out in triplicate, three times.

Effect of Ptx on cell viability: SKOV3-Red, SKOV3-Green and SKOV3-M cells were seeded in 96-well plates at a density of 10’000 cells/well. 12h later, cells were treated with 0, 10, 25, 50 and 100 nM of Ptx for 24h. Culture medium was then replaced by 100μl of MTT reagent. Plates were analyzed after 2 hours using a microplate reader at 540 and 690 nm. These experiments were carried out in triplicate, three times.

### 8.- Invasion assay

Cell invasion assay was performed in an invasion chamber as described elsewhere [24]. Briefly, 15 × 10^3^ cells (SKOV3-Red, SKOV3-Green and SKOV3-M) in 100 μl of RPMI supplemented with 10% FBS and 1% gentamicin were added to the upper compartment of the transwell chambers. RPMI supplemented with 20% FBS and 1% gentamicin (400 μl) was added in the lower chamber for 48 h at 37 °C in a CO_2_ (5%) incubator. After incubation, viable cells that invaded collagen were stained with crystal violet and measurement was performed at 540 nm. This assay was repeated three times and each experiment was run in triplicate. Data were normalized by proliferation values (MTT assay) and expressed as AU (invasion)/ AU (MTT).

### 9.- Tumor development on chick chorioallantoic membrane (CAM)

Tumor development on CAM was performed as previously described [25]. Briefly, fertilized eggs (animal facility of the University of Geneva, Geneva, Switzerland) were incubated at 38ºC with 80% relative humidity and periodic rotation. On egg development day (EDD) 4, the eggs were drilled at their narrow apex and the hole was closed with adhesive tape. The eggs were then incubated at 38ºC with 80% relative humidity without rotation. On EDD8 the hole in the eggshell was enlarged to allow the access to the CAM. After gently scratching of the membrane with a needle tip, SKOV3-Red, SKOV3-Green or SKOV3-M cells suspension (2×10^6^ cells in 30 μl of geltrex, ThermoFischer Scientific) were inoculated and the hole was hermetically covered with Parafilm^®^. Eggs were returned to the incubator to allow tumor growth. Tumor growth was then monitored at EDD10.5 and 13.5 using a Wild Heerbrugg M3Z microscope at 10x magnification with a Lumenera INFINITY2-1 CDD camera with Infinity Capture Software.

### 10.- Polyploidy index / Fusion index

#### Non-transfected cells

To visualize syncytia, SKOV3 and COV318 cells were fixed in paraformaldehyde (PFA) 4% at 4°C for 10 min, washed three times in PBS and stained with Hematoxylin (Sigma, Darmstadt, Germany) for 1 min. Cells were then washed twice with warmed tap water before bright-field imaging (EVOS, Life Technologies, Carlsbad, USA).

#### SKOV3-M

To quantify fusion index in SKOV3-M cells, cells were seeded in Nunc Lab-Tek II Chamber slides (Life Technologies, Carlsbad, USA) and processed for treatment. Cells were then fixed in PFA 4% at 4°C for 10 min, washed three times in PBS and incubated in PBS-3% bovine serum albumin (BSA) for 30 min to eliminate non-specific binding. Cells were then stained with Alexa Fluor 647 phalloidin (1:40 dilution, from Invitrogen, Thermo scientific) for 20 min at room temperature and rinsed three times in PBS. Slides were then mounted with Vectashield with DAPI (Vector Laboratories, Burlingame, USA) and sealed prior imaging with confocal fluorescent microscope (LSM 800 Airyscan, Zeiss, Iéna, Germany). Images were processed using ImageJ freeware.

The polyploidy or fusion index expressed in percent was calculated as follows: [(N-S)/T]×100, where N equals the number of nuclei in syncytia, S equals the number of syncytia and T equals the total number of nuclei counted [26]. This index was calculated for three independent experiments, run in triplicate.

### 11.- Fluorescent microscopy

To visualize syncytia in SKOV3-Green, SKOV3-Red and SKOV3-M cells, cells were seeded in μ-Slide 8 Well (Ibidi, Madison, USA) and processed for treatment. Cells were then fixed in PFA 4% at 4°C for 10 min, washed three times in PBS and incubated in PBS-3% BSA for 30 min to eliminate non-specific binding. Cells were then stained with Alexa Fluor 647 phalloidin (1:40 dilution, from Invitrogen, Thermo scientific) for 20 min at room temperature, rinsed three times in PBS and stained with DAPI for 10 min. Cells were then washed three times in PBS before imaging with fluorescent microscope (EVOS FL, Life Technologies, Carlsbad, USA).

### 12.- Western blot

Whole cell extracts (40 μg of proteins) were fractionated by SDS-Page 10% and transferred to nitrocellulose membrane for immunoblot analysis using rabbit anti-GRP78 antibodies (GL-19, 1:5000 dilution from Sigma), anti-PERK antibodies (CS56833, 1:1000 dilution from Cell Signaling), mouse anti-CHOP antibodies (MA1-250, 1:500 dilution from Invitrogen) and mouse anti-GAPDH antibodies (1:60 000 dilution from Millipore).

### 13- Microarray

Microarray-based transcriptome profiling was performed at the iGE3 Genomics platform of faculty of medicine of the University of Geneva (https://ige3.genomics.unige.ch). Briefly, 100 ng of total RNA were extracted from SKOV3-Red, SKOV3-Green and SKOV3-M at two different passages and were used as input for the preparation of single-strand cDNA using the WT PLUS reagent kit from Thermofisher Scientific. Targets were then fragmented and labeled with the Affymetrix GeneChip WT Terminal Labeling Kit and hybridized on Human Clariom S arrays according to manufacturer’s recommendations.

The obtained data were analysed with the Transcription Analysis Console (TAC) software (ThermoFisher Scientific), selecting the genes which expression was modified at least 2-fold and which p-value was <0.05.

### 14.- Statistical analysis

Data were represented as means ± SEM for at least 3 different samples. Statistical differences between samples were assessed by the Student’s t test. P-value <0.05 was considered as significant. GraphPrism software was used to perform the different statistical analysis.

## 3. Results

### 3.1. PGCCs in ovarian cancer cell lines originate, at least partially, from cell fusion

PGCCs origin remains elusive, since both endoreplication and cell fusion are potential mechanisms that can trigger the emergence of these cells. In order to confirm the fusion origin of PGCCs in ovarian cancer, we used two different ovarian cancer cell lines, SKOV3 and COV318. Our first approach consisted in mixing and growing two populations of the same cell line, which had previously been stained with green or far red cytoplasmic dyes as previously described by [27]. After double-positive flow cytometry sorting, we obtained a heterogeneous population of cells displaying both cell dyes. Unexpectedly, the outcome of the cell sorting was not formed mostly by fused cells that mixed their cytoplasm; instead, it was constituted by a mix of polynucleated cells and mononucleated cells that exchanged vesicles with cells stained with the opposite cytoplasmic marker (Figure 1A). This result evidenced that this approach previously used in other publications is not valid in these cells since vesicle exchange happened and interfered with the results.

**Figure 1.**
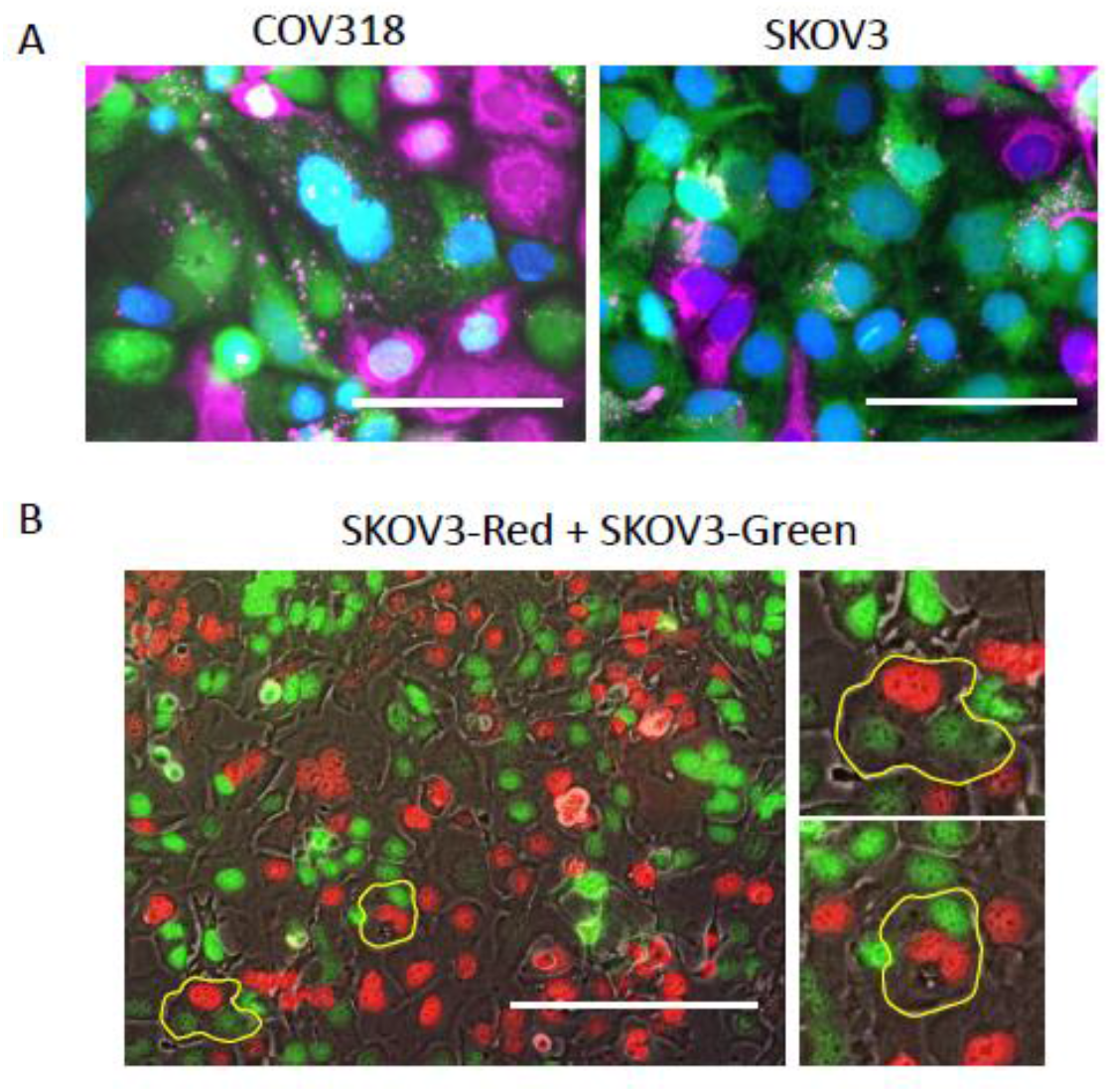
Formation of ovarian cancer PGCCs by cell fusion. **A-** Ovarian cancer cells (COV318 and SKOV3) double positive for green and far-red cytoplasmic dyes were sorted by flow cytometry following induction of cell fusion with 300 μM CoCl_2_ for 48h. Scale bars represent 100 μm. **B-** SKOV3 cells were transfected with plasmids encoding for red (SKOV3-Red) and green (SKOV3-Green) nuclear fluorescent proteins and mixed in co-culture. Under regular culture conditions, cells spontaneously formed PGCCs by cell fusion, visualized by polynucleated cells with both red and green nuclei (circled in yellow). Scale bars represent 200 μm (with two high magnifications of selected portions at the bottom).

With the purpose of eliminating the false positive results obtained, we decided to mark permanently the nuclei of SKOV3 cells by transfecting them with plasmids expressing a nuclear protein tagged with red or green fluorescent proteins and different antibiotic resistance genes. Following antibiotic selection of transfected cells, we obtained homogenously nuclei marked cell lines, for now on referred as SKOV3-Red and SKOV3-Green cell lines. After mixing and growing the two populations of cells, polynucleated cells with red and green nuclei could be observed (Figure 1B), demonstrating that spontaneous cell fusion process is, at least partially, involved in the generation the PGCCs.

### 3.2. PGCCs nuclei are able to fuse

In order to isolate PGCCs formed by cell fusion, meaning that they contained green and red nuclei, we took advantage of the transfected SKOV3 cells described above. Indeed, fused cells containing both types of nuclei would be able to survive a double antibiotic selection, while the unfused parental cell lines and the same type of nuclei fused cells would die under this condition. Strikingly, after three weeks under double antibiotic selection, we observed that SKOV3 fused cells were no longer polynucleated and that their single nucleus was positive for both fluororescent proteins. In fact, the overlay of both red and green signals in the same nucleus was visible in yellow under fluorescent microscope (Figure 2), resulting from nuclei fusion. We then decided to characterize this new cell line, referred from now on as SKOV3-M.

**Figure 2:**
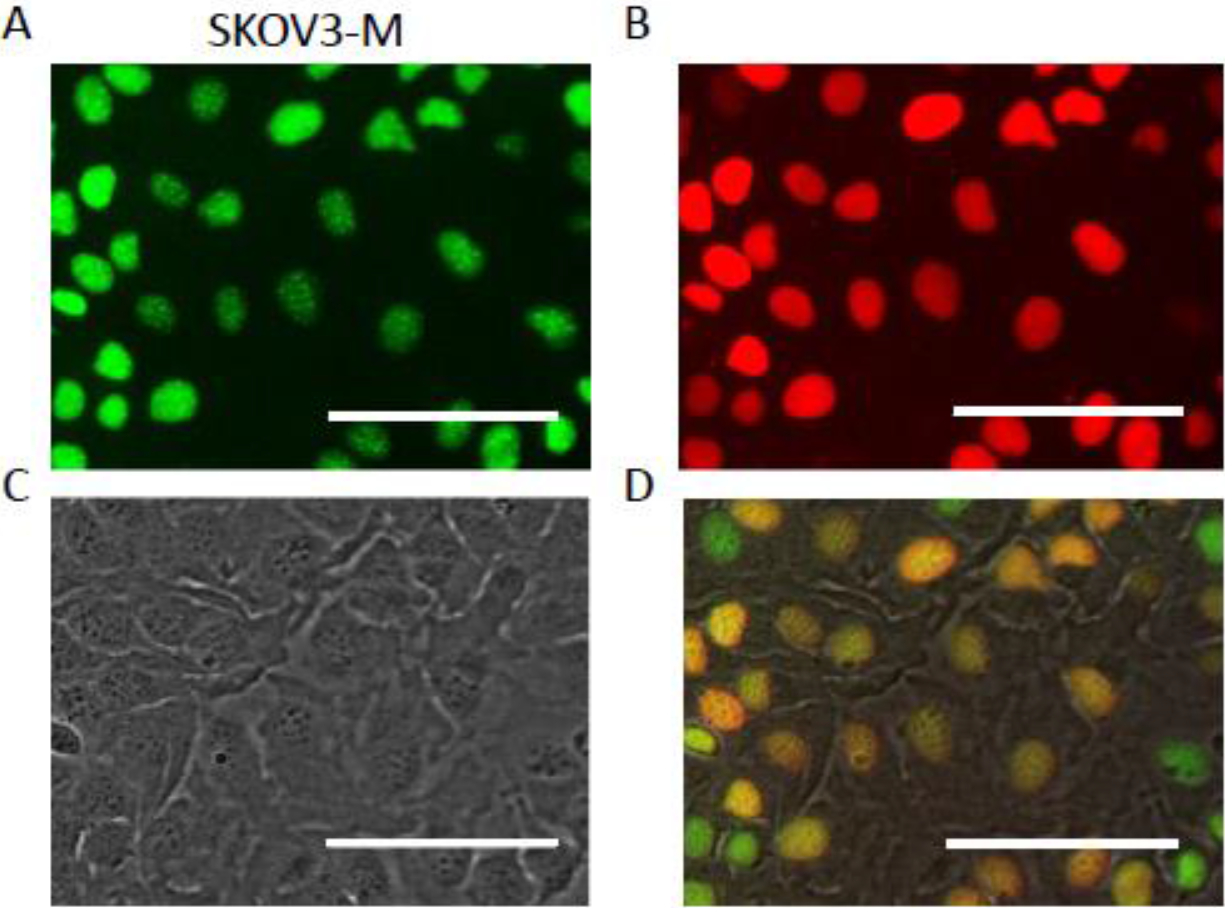
Ovarian cancer PGCCs can undergo nuclear fusion. SKOV3 cells were transfected with plasmids encoding for red (SKOV3-Red) and green (SKOV3-Green) nuclear fluorescent proteins, associated with two different antibiotic resistance genes. SKOV3-Red and SKOV3-Green were mixed in co-culture with double antibiotic selection (100 μg/ml G418 and 0.2 μg/ml puromycin). After three weeks of double selection, all cells displayed single nucleus double positive for both green **(A)** and red **(B)** signal, proving previous fusion of red and green nuclei. This new cell line is thereafter mentioned as SKOV3-M. **C-** Phase contrast. **D-** Overlay. Scale bars represent 100 μm.

#### 3.2.1. Transcriptomic analysis

To characterize the SKOV3-M cell line, we first performed a transcriptomic analysis, to get a hint of the possible modified processes in these cells. We compared the expression levels of RNAs in SKOV3-M cells with the ones in SKOV3-Green and SKOV3-Red cells. We obtained 312 genes which expression was at least 2-fold changed; 173 genes were upregulated and 139 genes were downregulated in SKOV3-M cells when compared with the unfused parental cells. None of them was related to cell cycle, suggesting that SKOV3-M cells were subjected to similar cell cycle regulation than the parental cell lines. However, some of these genes were related to invasiveness, cell motility and cell adhesion and were listed in Table 1.

**Table 1.** Cell migration, cell motility, invasion, cell adhesion and locomotion-related genes which expression is at least 2-fold modified and which p-value is <0.05 in SKOV3-M cells compared to SKOV3-Green and SKOV3-Red cells.

| *Gene symbol* | *Fold change* | *P-value* | *Function* |
| --- | --- | --- | --- |
| IL1R2 | 11.03 | 5.98<sup>E</sup>-06 | Cell migration |
| MMP2 | 8.13 | 0.002 | Invasion |
| MXRA5 | 7.09 | 2.79E-05 | Invasion |
| VEGFC | 3.57 | 0.0108 | Invasion |
| AFAP1L2 | 3.1 | 0.0012 | Cell motility |
| EFNB2 | 2.71 | 2.95E-05 | Invasion |
| LRRC17 | 2.5 | 0.0032 | Locomotion |
| ADGRE5 | 2.49 | 1.96E-05 | Cell adhesion |
| ELFN2 | 2.34 | 0.0002 | Cell migration |
| FBLN5 | -2.02 | 0.0043 | Cell adhesion |
| BCAM | -2.15 | 0.0012 | Cell adhesion |
| CDHR1 | -2.54 | 6.17E-05 | Cell adhesion |
| ITGB4 | -2.56 | 0.0197 | Cell adhesion |
| IL7R | -3 | 0.0064 | Invasion |
| **ADAMTSL4** | -3.33 | 0.0042 | Cell adhesion |
| **EPCAM** | -3.99 | 1.31E-06 | Cell adhesion |
| **IGFBP3** | -5.18 | 0.0007 | Cell migration |
| **S1PR1** | -6.54 | 1.65E-07 | Cell migration |

#### 3.2.2. Proliferative capacities of SKOV3-M cells

Given the fact that cell cycle related genes expression was not modified in the transcriptome analysis, and SKOV3-M cells were no longer polynucleated cells, we decided to analyse the cell cycle in these cells. We observed a similar G0-G1 and G2 phase cell distribution between SKOV3-M and its parental cell lines, SKOV3-Green and SKOV3-Red cells (Figure 3A, B). Interestingly, the Hoechst staining, which marked the DNA, was more intense in SKOV3-M cells (Figure 3A), confirming a higher DNA content in the fused cells than in the parental ones. Additionally, we decided to study the proliferative capacities of SKOV3-M cells by staining the cells with CellTrace™ far red. The division profile of stained cells was analysed by flow cytometry during five consecutive days. At the end, SKOV3-M and its parental cell lines had proliferated and diluted CellTrace™ far red fluorescence in exactly the same way (Figure 3C, D). Given the similar proliferative capacities between SKOV3-M and the parental cell lines measured by flow cytometry, we tested the capacity of these cells to resist to the antimitotic drug Paclitaxel (Ptx). Measurement of cell viability after 24h treatment with increasing doses of Ptx showed that SKOV3-M cells resisted in a similar manner to Ptx than the parental cell lines (Figure 3E) suggesting that the capacity of these cells to divide again renders them sensitive to Ptx.

**Figure 3:**
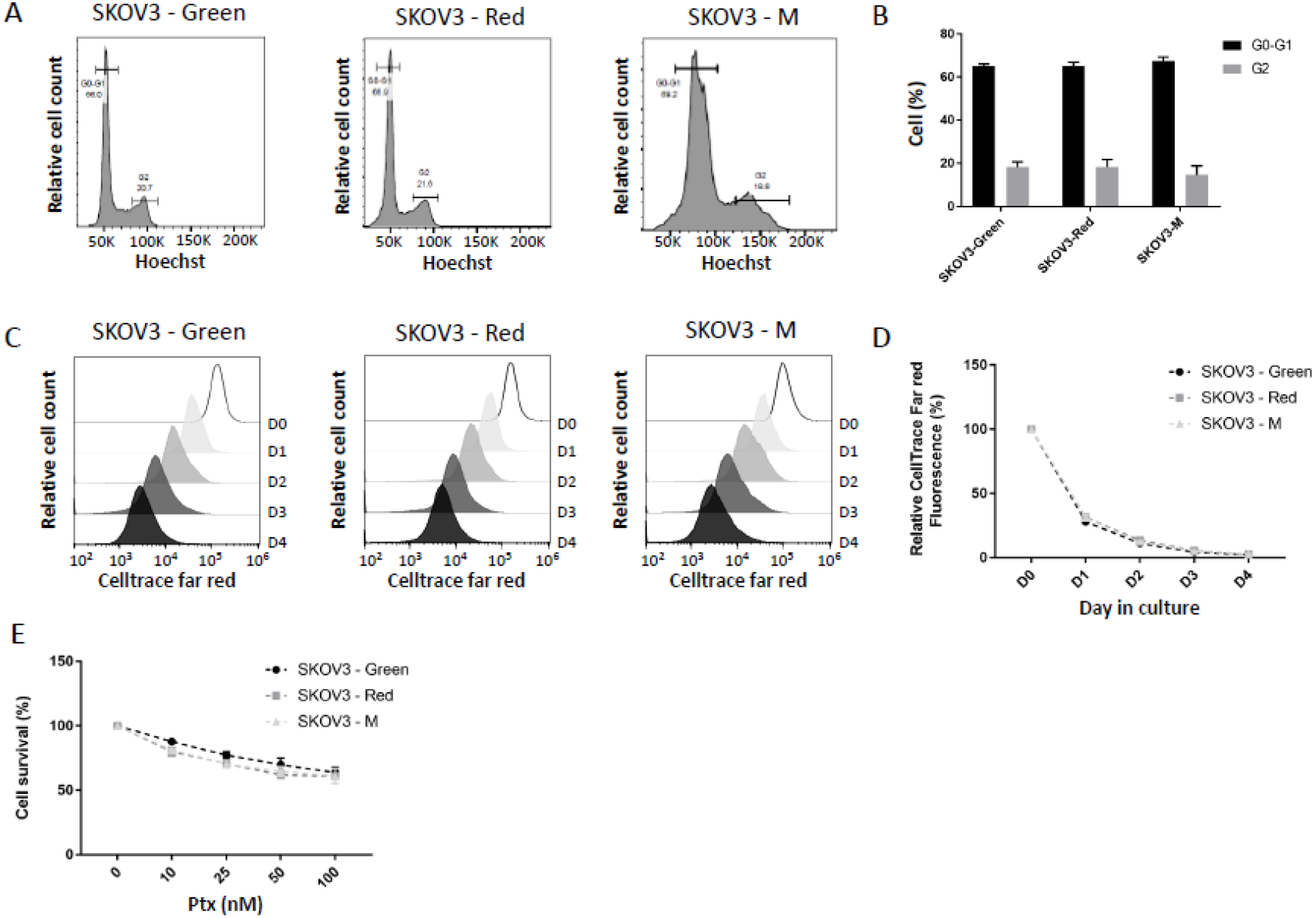
SKOV3-M cells display normal proliferation capacities and cell cycling. **A and B-** Cell cycle distribution was analyzed in SKOV3-Red, SKOV3-Green and SKOV3-M cells by flow cytometry following Hoechst staining. Doublets cells have been removed from the analysis based on Hoescht W and Hoescht A signals. **A-** Cell distribution depending on Hoechst signal intensity. **B-** Percentages of cells in G0-G1 and in G2 phases. **C and D-** Proliferation capacities of SKOV3-Red, SKOV3-Green and SKOV3-M cells were assessed by analyzing CellTrace™ far red cytoplasmic signal dilution by flow cytometry. Cells were stained on day 0 (D0) and seeded in 10 cm culture dishes (5 × 10^5^ cells/dish). Far red signal was analyzed for five consecutive days (D0 to D4). Doublets cells have been removed from the analysis based on SSC W and SSC A signals. **C-** Evolution of far red signal intensity from D0 to D4. **D-** Evolution of far red signal dilution expressed as percentage of mean signal intensity ± SEM measured at D0. **E-** SKOV3-Red, SKOV3-Green and SKOV3-M cells were treated with increasing doses of Paclitaxel (Ptx). Cell survival was assessed by MTT assay after 24h of treatment. Results are expressed as mean percentages ± SEM of optical density measured in control cells.

#### 3.2.3. Invasive properties of SKOV3-M

Transcriptomic analysis displayed modified RNA expression of invasiveness related genes (Table 1). Increased invasive properties would confer SKOV3-M cells the advantage to propagate cancer. To test this hypothesis, the three cell lines SKOV3-Green, SKOV3-Red and SKOV3-M were seeded in Boyden chamber. The capacity of SKOV3-M to invade collagen was increased compared to its parental cell lines (Figure 4A).

**Figure 4:**
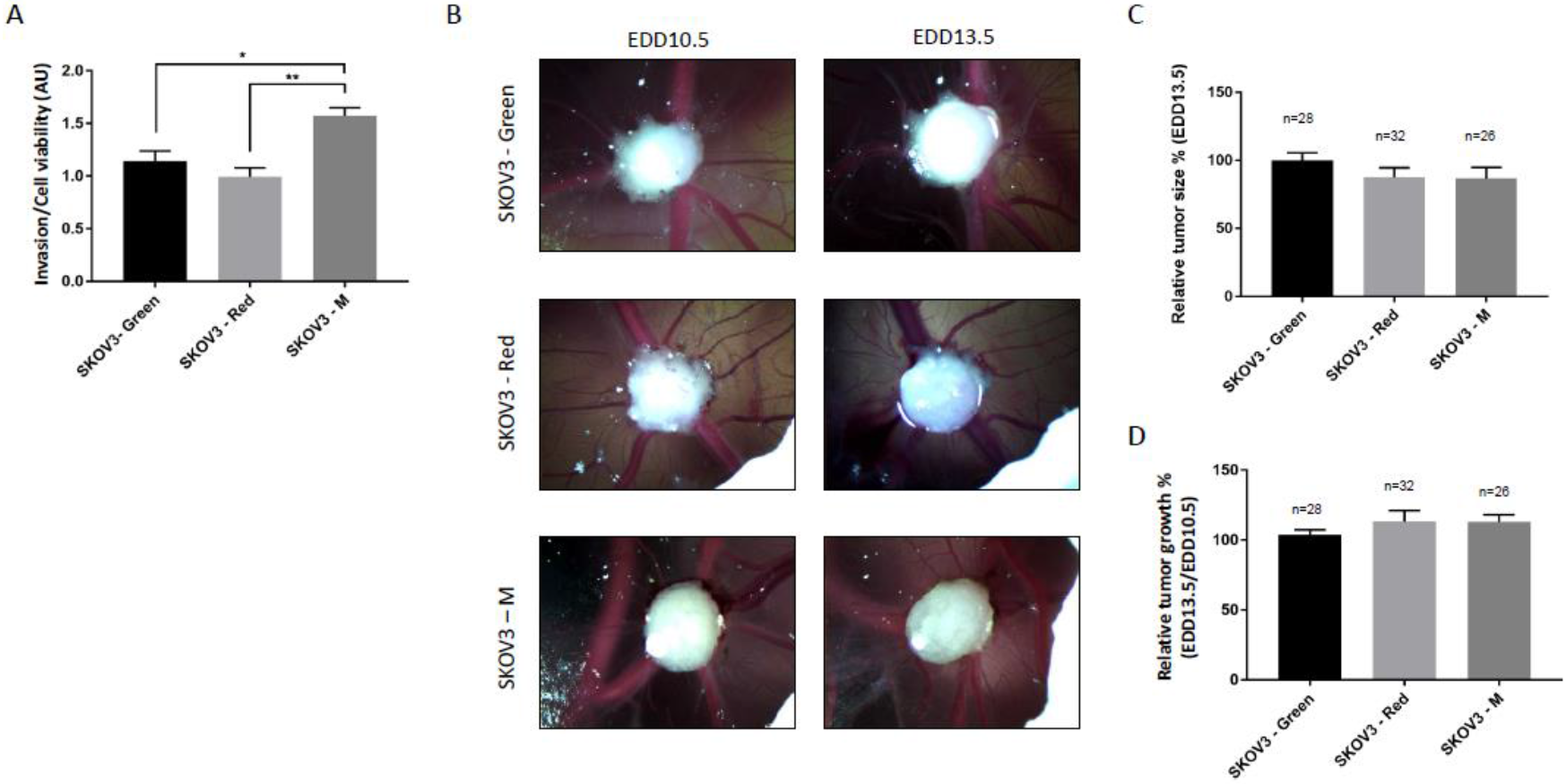
SKOV3-M cells display enhanced invasive capacity and unaffected tumorigenic ability. **A-** Invasive capacity of SKOV3-Red, SKOV3-Green and SKOV3-M cells was assessed by the Boyden chamber assay. Cell viability was assessed by MTT assay. Results are expressed as invasion relative to cell viability ± SEM. AU: arbitrary unit, * p<0.05, ** p<0.01. **B to D-** Tumorigenic capacity of SKOV3-Red, SKOV3-Green and SKOV3-M cells was assessed using chick chorioallantoic membrane (CAM) model. Cells were inoculated on egg development day (EDD) 8, and tumor size was measured at EDD10.5 and EDD13.5. **B-** Images of the tumors growing in CAM at EDD10.5 and EDD13.5 **C-** Tumor size measured at EDD13.5 for SKOV3-Red, SKOV3-Green and SKOV3-M relative to tumor size formed with SKOV3-Green cells. **D-** Tumor growth (tumor size at EDD13.5 relative to tumor size at EDD10.5). Results are expressed as mean ± SEM.

#### 3.2.4. Tumor forming abilities of PGCCs

We then investigated *in ovo* the capacity of SKOV3 cell lines to form tumors. We inoculated 2×10^6^ SKOV3-M, SKOV3-Green or SKOV3-Red cells on chick chorioallantoic membrane (CAM) on egg development day eight (EDD8) and monitored tumor growth at EDD10.5 and EDD13.5 (Figure 4B). The tumor size (Figure 4C) and tumor growth (Figure 4D) were not different between SKOV3-M and the parental cell lines, suggesting equivalent tumor forming abilities.

### 3.3. - ERS increases cancer cell fusion

The different processes that are implicated in cancer cell fusion are not deeply investigated. It was shown that chemical hypoxia, achieved with CoCl_2_ treatment, or physiological hypoxia, achieved by 0.1% oxygen cell culture, induced ovarian PGCCs formation [9,13–15]. The hypoxic conditions, which are known to exist in the tumor environment, are capable of eliciting the UPR. Interestingly, we have shown that GRP78, master regulator of the UPR, is involved in physiological cell fusion leading to formation of syncytiotrophoblast [22]. In order to test the UPR involvement in ovarian cancer cell fusion, we treated SKOV3 and COV318 cells with ERS inducers. The three ERS inducers –Thapsigargin (THA), Tunicamycin (TUN) and HA15– managed to increase GRP78 and CHOP expression (Figure 5A,B). The resulting UPR activation led to a significant increase in cell fusion in all of the tested conditions (Figure 5C,D). For further investigation, we decided to keep on working with HA15 as an activator of the UPR since THA and TUN had slight cytotoxic effects (data not shown). In order to demonstrate that the UPR inducer has an effect on cell fusion, we mixed SKOV3-Green and SKOV3-Red cells and treated them with HA15 before quantifying the cell fusion rate. Using confocal microscopy, we observed an increased amount of polynucleated cells containing red and green nuclei. Based on these images, the fusion index was measured (Figure 5E, F), confirming that ERS induced cell fusion.

**Figure 5:**
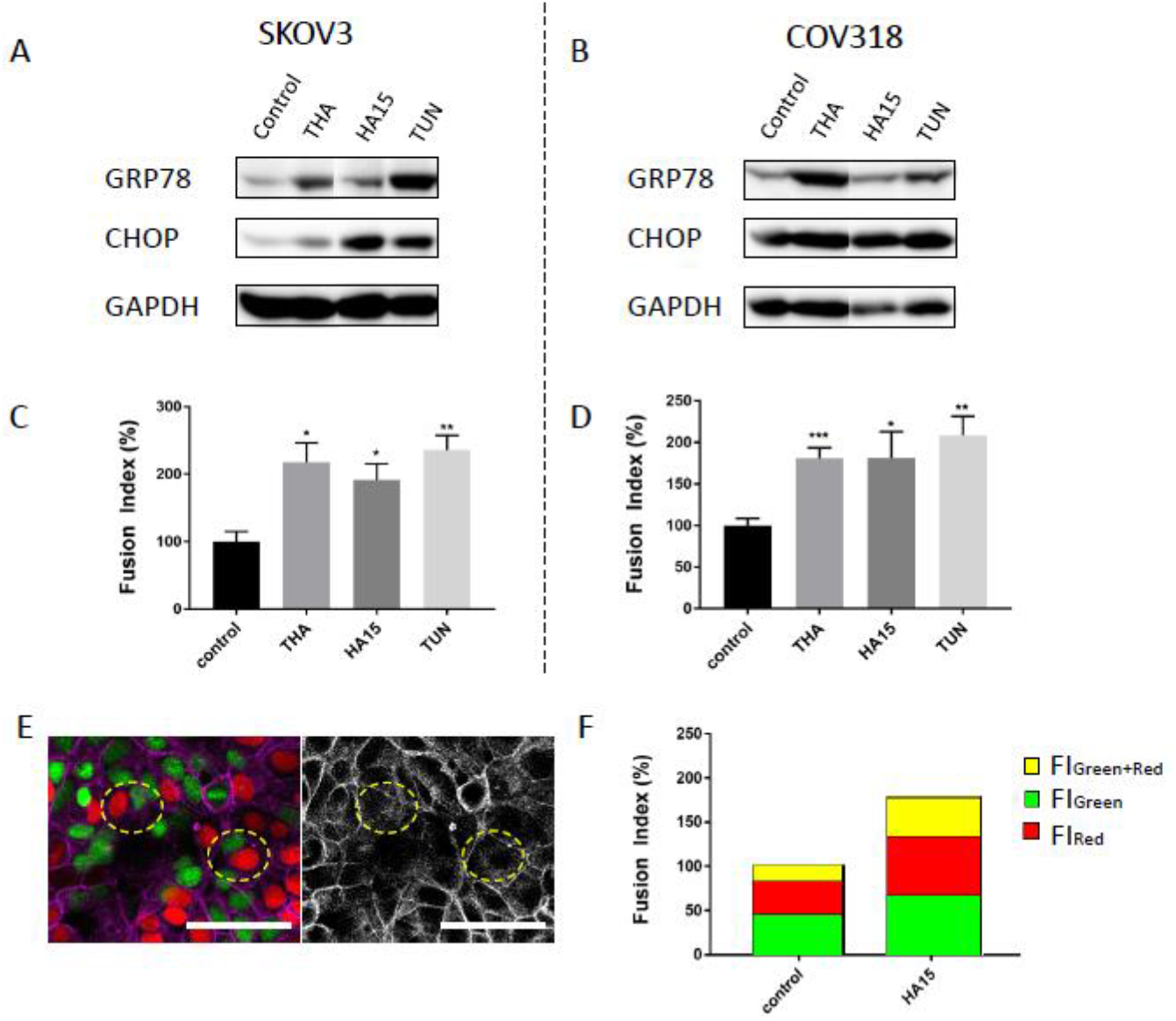
ERS induction enhances cell fusion in ovarian cancer cells. **A to D-** SKOV3 (**A and C**) and COV318 (**B and D**) cells were treated with three different ERS inducers, 100nM Thapsigargin (THA), 1 μM HA15 or 1 μg/mL Tunicamycin (TUN) for 48h. Non-treated cells were used as control condition. **A and B-** Protein levels for GRP78, CHOP and GAPDH were assessed by Western blotting. **C and D-** Fusion index was calculated for the different treatments and expressed as percentages relative to control condition. Results are expressed as mean ± SEM. * p<0.05, ** p<0.01, *** p<0.001. **E and F-** Co-culture of SKOV3-Green and SKOV3-Red cells was treated with 1 μM HA15 for 48h. **E-** Presence of polynucleated cells with both red and green nuclei was detected by confocal microscopy. Scale bars represent 100 μm. **F-** Fusion index was calculated separately for green polynucleated cells (Fl_Green_), red polynucleated cells (Fl_Red_) and red/green polynucleated cells (Fl_Green+Red_). Results are expressed as mean percentages relative to fusion index calculated for control condition.

We then used molecules known to inhibit UPR activation. Since no specific inhibitors of ATF6 activation were available, we used 4-(2-aminoethyl) benzenesulfonyl fluoride hydrochloride (AEBSF, which blocks S1P and S2P in the Golgi apparatus, preventing ATF6 cleavage) and Melatonin (Mel) [28] to inhibit ATF6 pathway activation. STF-083010 (STF) and 4μ8C were used to inhibit IRE1*α* Rnase. Finally, GSK2656157 (GSK, an ATP competitive inhibitor), and Salubrinal (SAL, an inhibitor of EIF2*α* dephosphorylation) were used to modulate PERK pathway activation. When SKOV3 cells were simultaneously treated with THA and UPR activation inhibitors, we observed a general lower cell fusion rate compared to THA alone (Figure 6A). However, the only significant reduction was achieved when treating the cells with the ATF6 pathway inhibitors. Differently, treatment of SKOV3 cells simultaneously with TUN and UPR inhibitors showed more homogeneously significant reduction in cell fusion rate (Figure 6B). In the same way, COV318 treatment with THA or TUN and the different UPR inhibitors resulted in a significant cell fusion rate decrease, except for the combination of TUN + AEBSF (Figure 6C,D). Together, these results tend to confirm the role of UPR, probably through the activation of the three UPR sensors ATF6, IRE1*α* and PERK, in ovarian cancer cell-cell fusion.

**Figure 6:**
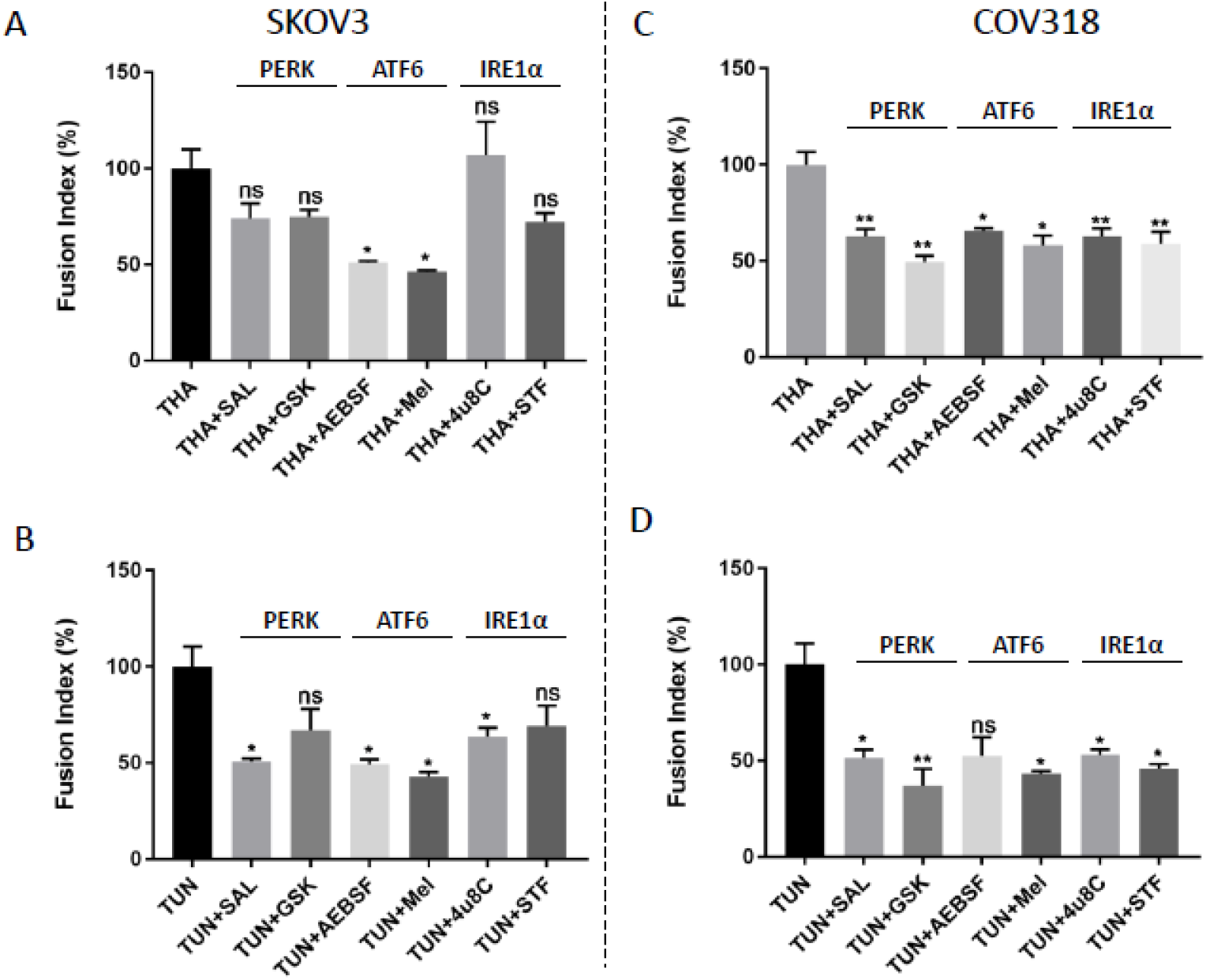
UPR pathways inhibition differentially affects cell fusion in SKOV3 and COV318 cell lines. SKOV3 (**A and B**) and COV318 (**C and D**) cells were treated either with 100nM Thapsigargin (THA) (**A and C**) or 1 μg/mL Tunicamycin (TUN) (**B and D**), combined with inhibitors of the three UPR pathways: 17.5 μM Salubrinal (SAL) or 0.3 μM GSK2656157 (GSK) for PERK pathway inhibition, 200 μM 4-(2-aminoethyl) benzenesulfonyl fluoride hydrochloride (AEBSF) or 1 mM Melatonin (Mel) for ATF6 pathway inhibition, and 8 μM 4μ8C or 2 μM STF-083010 (STF) for IRE1*α* pathway inhibition, for 48h. Fusion index was expressed as percentages relative to fusion index calculated for either THA or TUN condition. Results are expressed as mean ± SEM. ns: non significant, * p<0.05, ** p<0.01.

PERK is known to promote eIF2*α* phosphorylation, and thus the translation of ATF4 [review at [18]]. ATF4 is one of the transcription factor family members of cAMP response element binding (CREB) and was shown to promote expression of fusogen, syncytin 2, and thus cell fusion in a choriocarcinoma cell line [29]. We thus decided to specifically investigate the implication of PERK pathway in ovarian cancer cell fusion.In order to selectively silence PERK UPR-branch, we then used siRNA gene-silencing method. The impact of PERK silencing in cell fusion was measured by treating SKOV3 and COV318 cells with the UPR activators at the same time that PERK was silenced. The knockdown of PERK was preliminary validated by checking the PERK protein levels by Western blot. In SKOV3 cells, PERK protein expression was efficiently reduced when the silencing was performed in control or cells treated with ERS inducers (Figure 7A). PERK silencing in SKOV3 cells treated with UPR inducers resulted in a decreased protein expression of GRP78, suggesting the expected lower ERS activation (Figure 7A). PERK knockdown and its linked ERS inhibition significantly decreased the fusion rate in SKOV3 cells (Figure 7B). The same experiment was performed in COV318, firstly showing the efficient PERK inhibition and consequent UPR activation or inhibition when measuring PERK, GRP78 and CHOP expression by Western blot (Figure 7C). COV318 cells showed as well a homogeneous response to PERK expression down regulation, displaying significant cell fusion inhibition when the UPR was simultaneously activated with either THA, TUN or HA15 (Figure 7D). Interestingly, PERK silencing in control SKOV3 and COV318 cells caused an increased expression of GRP78 (Figure 7A,C) suggesting that the decrease in PERK expression may lead to a burst of UPR activity. The UPR activation resulting from PERK knockdown increased significantly cell fusion in both SKOV3 and COV318 cells (Figure 7B, D), which reinforces the UPR involvement in cell fusion.

**Figure 7:**
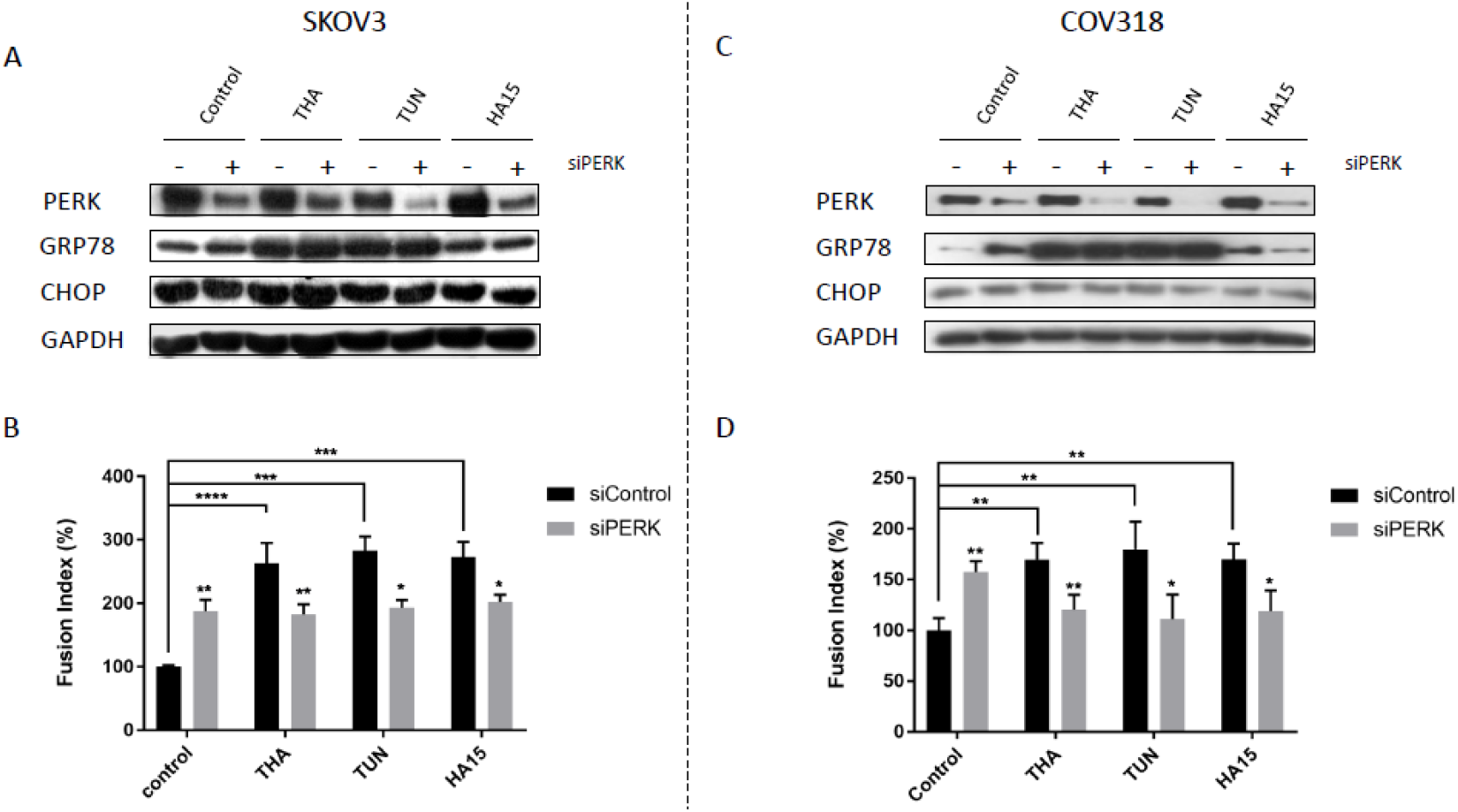
PERK silencing differentially reduces cell fusion in SKOV3 and COV318 cell lines. SKOV3 (**A and B**) and COV318 (**C and D**) cells were transfected either with 10 nM PERK siRNA (siPERK) or 10 nM control siRNA (siControl) and then treated 24h later with three different ERS inducers, 100nM Thapsigargin (THA), 1 μM HA15 or 1 μg/mL Tunicamycin (TUN) for 48h. Cells treated with DMSO were used as control condition. **A and C-** Protein levels for PERK, GRP78, CHOP and GAPDH were assessed by Western blotting. **B and D-** Fusion index was calculated for the different treatments and expressed as percentages relative to fusion index calculated for control condition in siControl cells. Results are expressed as mean ± SEM. * p<0.05, ** p<0.01, *** p<0.001, **** p<0.0001

## 4. Discussion

The PGCCs formation has been studied in several cancer types, among which colon, breast and ovarian cancer [9,13–15]. This process is induced by DNA damaging agents and could be linked to chemotherapeutic resistance acquisition [9,13,14,30]. It could be also incremented by chemical hypoxia treatment (CoCl_2_) or hypoxic culture conditions *in vitro* [9,13–15]. The mechanism of PGCCs formation has always been controversial since it could be explained either by a defect in cytokinesis leading to endoreplication, by cell fusion events or by a combination of both mechanisms [8,9,12]. Under chemical hypoxia, it was shown that at least 10 to 20% of ovarian PGCCs were formed by cell fusion [9]. The characterization of the PGCCs formed under these conditions displayed a modification in their division process, changing from mitosis to bursting and budding [9].The PGCCs generated daughter cells via asymmetric division were able to form spheroids and expressed several cancer stem cell markers [9]. Moreoever, they were more tumorigenic than regular-sized cancer cells in the nude mice [9]. In the present paper, using plasmids expressing nuclear proteins tagged with green or red fluorochrome, we generated PGCCs from SKOV3 cells and confirmed that these giant polynucleated cells were at least partially derived from cell fusion events. Moreover, we described for the first time, that under double antibiotic selection pressure, PGCCs nuclei were able to fuse leading to homogeneous mononucleated cell line that we called SKOV3-M which nuclei contain proteins with both red and green fluorescence. Despite higher DNA content, SKOV3-M cells recovered normal cell cycle and were able to proliferate in the same manner as their parental cell lines (SKOV3-Green and SKOV3-Red). This novel observation leads us to hypothesize that nuclear fusion may be a mechanism by which PGCCs, after surviving drugs treatment, could change their cell division back to mitosis. Transcriptomic analysis of the SKOV3-M cells showed that 312 genes were at least 2-fold modulated in comparison with their parental cells. Among them, we found genes implicated in cell adhesion, mobility and invasion. The validation of these results with invasion assays demonstrated increased invasive properties of SKOV3-M cells compared to their parental cells. This increased invasiveness renders the cells more aggressive, favoring the onset of metastases [31–33]. The proliferation rate of these fusion-derived cells, their capacity to form tumors, which we measured *in ovo* on CAM model, and their invasive properties, could explain the high cancer relapse rates that ovarian cancer patients suffer. However, it remains the question whether ovarian cancer cell fusion actually occurs *in vivo*.

Despite that PGCCs could generate more aggressive daughter cells via bursting, budding [9,34,35] or mitosis [36] after nuclei fusion, the mechanism and conditions initiating their formation have been poorly studied. The increment in ovarian PGCCs formation achieved under hypoxia *in vitro* pointed towards hypoxia as a possible cell fusion enhancer [9,13–15]. Ptx treatment was also shown to cause an induced prominence of PGCCs [[13] and data not shown]. A shared pathway downstream of low oxygen tension and Ptx treatment is the endoplasmic reticulum (ER) stress-mediated UPR pathway [18,37–44]. We thus hypothesized that UPR activation may result in cancer cell fusion enhancement and PGCCs formation. The activation of the UPR with different UPR activators effectively increased cell fusion and PGCCs formation with both SKOV3 and COV318 cell lines. In the opposite, the inhibition of the UPR pathways by inhibitors targeting the three UPR-branches led to a general tendency of a cell fusion decrease. To confirm the involvement of UPR activation in cell fusion and PGCCs formation, we then specifically down regulated PERK pathway activation by decreasing PERK expression in ovarian cancer cells. The decrease in PERK expression in ovarian cancer cells led to an increase in UPR activation, which could be explained by the amplification of the signals transduced by the other two UPR pathways, as previously described in myeloma cells by Michallet et al. [45]. Indeed, the different UPR pathways display an extensive cross-talk between them, compensating their modulation and showing their interdependence [46,47]. However, under ERS conditions, the decrease in PERK expression in ovarian cancer cells led to a significant decrease in UPR activation and PGCCs formation. Therefore, we can conclude that the UPR, through the activation of the UPR sensors ATF6, IRE1*α* and PERK, is involved in ovarian PGCCs formation. However, the contribution of each UPR pathway and their interdependence remain to be elucidated.

More and more evidences showed that the UPR is involved in drug resistance in different cancer cells [40–42,44]. However, the implication of the UPR in chemotherapy resistance is complex and not fully understood. In this study, we showed that decreasing UPR activation in ovarian cancer cells could decrease ovarian cancer cell fusion and PGCCs formation. Avoiding the PGCCs formation may reduce cancer relapses and metastasis, while at the same time, the drug resistances generated by the UPR could be prevented [9,40,41,43]. A better understanding of the molecular mechanism controlling UPR-mediated PGCCs formation and drug resistance is highly needed to develop new combined therapeutic approaches with new drugs that specifically target the UPR sensors or downstream partners and conventional chemotherapy.

## Author Contributions

conceptualization, L.Y., D.B-R. and M.C.; methodology L.Y., D.B-R. and M.C.; formal analysis, L.Y., D.B-R. M.A. and M.C.; investigation, L.Y, D.B-R., C.W., P.R., M.A. and M.C.; resources, M.C. and P-Y.D.; writing—original draft preparation, L.Y., D.B-R. and M.C.; writing—review and editing, L.Y, D.B-R., P.R., M.A., P.P. and M.C.; visualization, L.Y. and D.B-R..; supervision, M.C.; project administration, M.C.; funding acquisition, M.C.

## Acknowledgments

We would like to thank the iGE3 Genomics and Flow Cytometry platforms’ staff for their technical support.

